# Accuracy of a markerless motion capture system for balance related quantities

**DOI:** 10.1101/2022.11.10.515951

**Authors:** Anaïs Chaumeil, Bhrigu Kumar Lahkar, Raphaël Dumas, Antoine Muller, Thomas Robert

## Abstract

**Background:** Balance studies usually focus on quantities describing the global body motion, such as the position of the whole-body centre of mass (CoM), its associated extrapolated centre of mass (XCoM) and the whole-body angular momentum (WBAM). Assessing such quantities using classical marker-based approach can be tedious and modify the participant’s behaviour. The recent development of markerless motion capture methods could bypass the issues related to the use of markers.

**Research question:** Can we use markerless motion capture systems to study quantities that are relevant for balance studies?

**Methods:** Sixteen young healthy participants performed four different motor tasks: walking at self-selected speed, balance loss, walking on a narrow beam and countermovement jumps. Their movements were recorded simultaneously by marker-based and markerless motion capture systems. Videos were processed using a commercial markerless pose estimation software, Theia3D. The position of their CoM was computed, and the associated XCoM and WBAM were derived. Bland-Altman analysis was performed and root mean square error and coefficient of determination were computed to compare the results obtained with marker-based and markerless methods across all participants and tasks.

**Results:** Bias remained of the magnitude of a few mm for CoM and XCoM position, and RMSE of CoM and XCoM was around 1 cm. Confidence interval for CoM and XCoM was under 2 cm except for one task in one direction. RMSE of the WBAM was less than 8% of the total amplitude in any direction, and bias was less than 1%.

**Significance:** Results suggest that the markerless motion capture system can be used in balance studies as the measured errors are in the range of the differences found between different models or populations in the literature. Nevertheless, one should be careful when assessing dynamic movements such as jumping, as they displayed the biggest errors.

**Highlights:**

- Markerless motion capture could bypass issues from classical marker-based approaches
- We compared balance related quantities computed from both approaches
- Mean differences were about 1cm on the position of the whole-body center of mass
- Obtained differences are acceptable for most applications

## Introduction

Balance studies have been conducted to understand the mechanisms that allow humans to maintain their balance in daily life activities such as walking [1], stair climbing [2] or rising from a chair [3]. These studies usually compute a range of biomechanical quantities which allow to describe the balance of an individual [4]. Whole body linear and angular momenta are widely studied [5–8] as they characterize the global movement of the body. While studying the position of the centre of mass (CoM) is common in balance studies [3], another interesting metric proposed by Hof [9] is the extrapolated centre of mass (XCoM): it is the CoM augmented by a proportion of its own velocity. Its position with respect to the border of the base of support (BoS) is an indicator of the balance of the participant [9]. These quantities are of fundamental importance for research in the balance area, and are classically estimated via a segmental approach, using the estimate of the positions, velocities and inertial parameters of the individual body segments.

Classically, balance related quantities are computed using marker-based motion capture, which is considered as the gold standard for motion analysis [10]. Although marker sets have been proposed to reduce the number of markers to be placed on the participant’s body [11], obtaining the segments and whole-body CoM still requires time and specific marker placement skills. Moreover, the associated experimental constraints limit the validity of the movement studied, as the participants are not in their usual environment, which can cause changes in behaviour [12].

During the last few years, the field of markerless motion capture systems based on video cameras has drastically expanded [13]. Several aspects of markerless motion analysis have yielded interest in the biomechanics community: ecological validity is enhanced as less strict experimental conditions are required, which allows for data acquisition in a real configuration. It also makes motion analysis more accessible, as it is possible to acquire and process data using low cost hardware such as regular smartphones or webcams and computers [14]. A large number of codes and software are now available, with different approaches and targeted applications [13,15]. One of them is Theia3D (Theia Markerless Inc., Kingston, Ontario, Canada, v2021.2) [16], a commercial software designed to study whole-body kinematics with minimal input from the user. Studies have shown that both upper [17] and lower [18,19] limbs kinematics have a good reliability [20] and are comparable to those computed using a marker-based system. The same conclusions hold for spatiotemporal gait parameters [19,21,22], both in standardized and clinical environment.

Considering the broad emerging literature in markerless motion analysis, few studies have focused on balance related quantities, and, when doing so, the authors mainly consider the CoM and its derivatives [23–28]. To the best of our knowledge, no study has been conducted regarding balance related quantities using Theia3D.

The goal of this study is to evaluate CoM, XCoM and whole-body angular momentum (WBAM) estimated with a markerless pose estimation system, Theia3D, in comparison to those obtained with a marker-based system for different motor tasks.

## Materials and methods

### Experimental session

Sixteen participants (9 men, 7 women) were recruited in this study, with mean age 25.1±3.0 years old, mean height 1.7±0.1 m and mean weight 68.4±13.5 kg. Participants had no history of musculoskeletal or balance problems. Prior to the experiment they signed an informed consent form. The study was approved by our institutional review board.

Marker-based optoelectronic and markerless video camera systems were set up for this experiment: 10 Qualisys Miqus M3 cameras recording at 300 Hz and 10 Qualisys Miqus Video recording at 60 Hz (1920 × 1088 pixels). The two systems were synchronized and spatially calibrated using Qualisys Track Manager (QTM – Qualisys AB, Sweden, v2021.1.2).

The participants wore minimal, tightly fitting clothing to allow for appropriate marker placement. They were equipped with a full-body marker set comprising 46 markers [29,30].

The participants performed a static trial (T pose) and four different motor tasks (Figure 1): walking at self-selected speed on a treadmill (**Walk**, between 7 and 20 cycles, Figure 1A), balance loss (**Lean**, lean forward until loss of balance requiring one or more recovery steps, 3 repetitions, Figure 1B), walking on a 2 m long, 2.3 cm wide and 5 cm high beam (**Beam**, 4 repetitions of walking the full length of the beam, Figure 1C) and countermovement jumps (**CMJS**, 3 repetitions, Figure 1D). These tasks were chosen as they were representative of different speeds (slow, normal, fast) and occurred in different planes of movement (sagittal and frontal planes).

**Figure 1:**
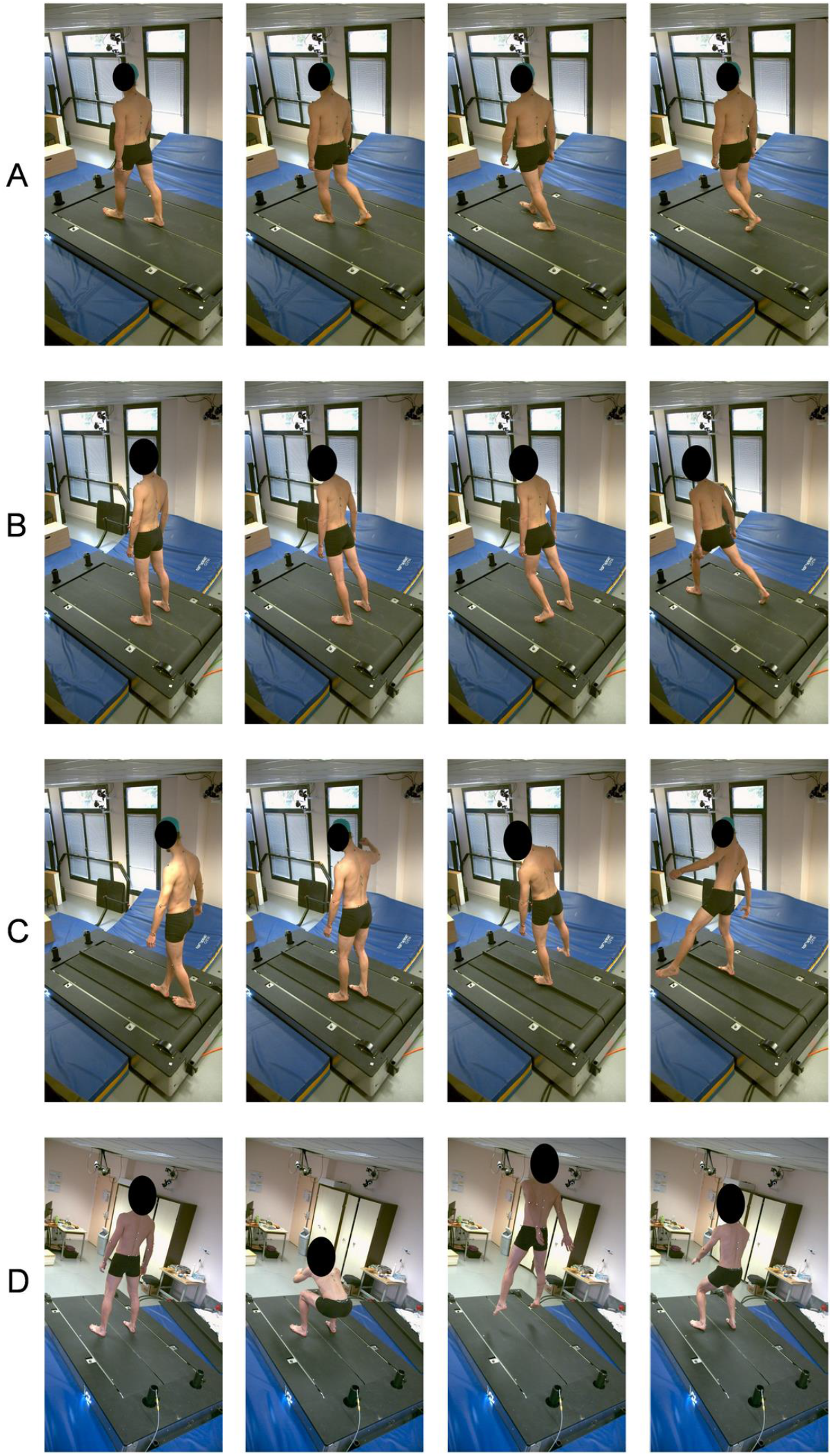
Snapshots of the tasks performed during the experimental session by a participant. A: Walk, B: Lean, C: Beam, D: CMJS.

### Data processing

#### Markerless data

Markerless data was processed with Theia3D, a deep learning-based commercial software. It uses deep learning techniques and inverse kinematics (IK) to estimate 3D pose of a subject of interest based on multi-camera video data and the associated calibration file [18]. The outputs of Theia3D were composed of a 17-segment kinematic model (head, thorax, upper arms, forearms, hands, pelvis, thighs, shanks, feet, toes) with 54 degrees of freedom (DoFs) and the pose matrices associated with each segment. These outputs were exported in Visual3D (C-motion, Germantown, USA, v2021.11.3) and body segment inertial parameters (BSIP – segment mass, position of segment CoM and moment of inertia) were added to the kinematic model according to Dumas and Wojtusch [29]. The markerless and marker-based models are both described in details in [30].

#### Marker-based data

Marker data was first processed in QTM, including labelling and gap filling of marker trajectories. Then, labelled and cleaned marker positions were imported in Visual3D where custom models were created based on Dumas and Wojtusch [29]. IK was finally performed using a multibody kinematics optimization [31]. The marker-based kinematic model was built in order to match as much as possible the markerless segments’ axes and kinematic model as described in Theia3D documentation. It has the same topology except that the toes are rigidly linked to the feet, and the same sex-specific BSIP.

#### Computing balance related quantities

For both marker-based and markerless data, the following quantities were exported from Visual3D following the IK step:

- the position of the CoM for each segment and the whole body;
- the linear velocity of the CoM of each segment;
- the angular velocity of each segment;
- the rotation matrix associated to the different segments.

Using MATLAB (MathWorks, USA), the positions and velocities of segments’ CoM and their angular velocities were filtered using a 4^th^ order Butterworth filter with a 7 Hz cut-off frequency. The velocity of the global CoM was obtained by differentiating the CoM and filtering the resulting signal with the same parameters. The XCoM was computed based on Hof [9] as:

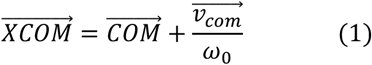

where 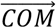 is the position of the whole-body’s 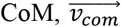 its velocity and 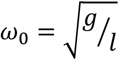 is a constant with *g* = 9.81*m*/*s*^2^ and *l* the height of the CoM of the participant during the static trial (T pose).

WBAM about the CoM was computed as:

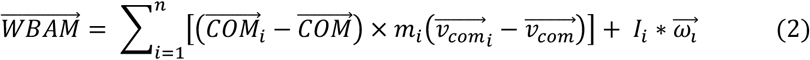

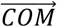 and 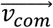 are respectively the position and velocity of the global 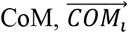 and 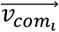 are respectively the position and velocity of the *i*^th^ segment’s CoM, *m*_*i*_ is its mass, *I*_*i*_ its inertial matrix expressed in the laboratory reference frame (computed using the inertias about the segment axes and the rotation matrix) and 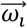 the angular velocity of the *i*^th^ segment with respect to the laboratory.

*WBAM* was made unitless as in [32] by dividing it by 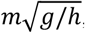, with *h* being the height of the participant and *m* it’s mass.

#### Statistical analysis

The level of similarity between marker-based and markerless results was assessed by computing Bland Alman bias and confidence interval (CI), as well as root mean square error (RMSE) and coefficient of determination (R^2^). The quantities of interest were CoM, XCoM and WBAM, the latter being expressed both unitless and as a percentage of the amplitude (computed as ±3 standard deviations across all motor tasks). All the statistical parameters were computed across all motor tasks and for each motor task separately, for all subjects.

## Results

Table 1 displays the statistical parameters bias, CI, RMSE and R^2^ for the CoM, XCoM and WBAM (expressed both as unitless and as a percentage of the total amplitude), computed across all tasks and for each task separately. Three directions are considered here: anterior-posterior (AP), medial-lateral (ML) and superior-inferior (SI). Bland Altman plots for CoM, XCoM and unitless WBAM can be found in Supplementary Materials.

**Table 1:** Results of the Bland-Altman analysis for each task and across all tasks, for all participants. For unitless WBAM, values have been multiplied by 10 000 (except R^2^) and displayed for easier reading.

|  |  | Walk |  |  | Lean |  |  | Beam |  |  | CMJS |  |  | All tasks |  |  |
| --- | --- | --- | --- | --- | --- | --- | --- | --- | --- | --- | --- | --- | --- | --- | --- | --- |
|  |  | AP | ML | SI | AP | ML | SI | AP | ML | SI | AP | ML | SI | AP | ML | SI |
| CoM (cm) | Bias | 0.068 | 0.099 | -0.53 | -0.59 | 0.11 | -0.17 | 0.17 | 0.018 | -0.24 | -0.30 | 0.087 | -0.16 | 0.012 | 0.065 | -0.27 |
|  | CI | 1.24 | 0.63 | 1.75 | 1.76 | 0.56 | 1.24 | 1.66 | 0.89 | 1.18 | 1.16 | 0.41 | 1.67 | 1.53 | 0.68 | 1.48 |
|  | RMSE | 0.64 | 0.31 | 0.86 | 1.03 | 0.29 | 0.68 | 0.80 | 0.44 | 0.72 | 0.74 | 0.24 | 0.86 | 0.79 | 0.37 | 0.77 |
|  | R<sup>2</sup> | 0.99 | 0.99 | 0.98 | 0.99 | 0.99 | 0.99 | 0.99 | 0.99 | 0.98 | 0.99 | 0.99 | 0.99 | 0.99 | 0.99 | 0.99 |
| XCoM (cm) | Bias | 0.044 | 0.058 | -0.51 | -0.62 | 0.079 | -0.27 | 0.20 | 0.019 | -0.27 | -0.27 | 0.062 | -0.10 | 0.0068 | 0.047 | -0.32 |
|  | CI | 1.74 | 1.075 | 1.88 | 1.94 | 0.78 | 1.45 | 1.80 | 1.18 | 1.46 | 1.83 | 0.89 | 2.76 | 1.87 | 1.061 | 1.69 |
|  | RMSE | 0.89 | 0.59 | 1.01 | 1.24 | 0.49 | 0.85 | 0.99 | 0.63 | 0.85 | 1.13 | 0.51 | 1.70 | 1.02 | 0.59 | 1.02 |
|  | R<sup>2</sup> | 0.99 | 0.99 | 0.98 | 0.99 | 0.98 | 0.99 | 0.99 | 0.98 | 0.98 | 0.99 | 0.97 | 0.99 | 0.99 | 0.98 | 0.99 |
| WBAM (unitless, x E-04 except R<sup>2</sup>) | Bias | -0.014 | 1.03 | -0.060 | 0.21 | 0.11 | -0.10 | -0.013 | 0.081 | 0.11 | 0.74 | -0.19 | -0.069 | 0.11 | 0.30 | 0.00016 |
|  | CI | 10.72 | 12.85 | 11.45 | 5.69 | 8.07 | 5.98 | 10.47 | 11.68 | 10.05 | 12.57 | 17.53 | 9.11 | 10.02 | 12.04 | 9.64 |
|  | RMSE | 5.46 | 6.75 | 6.37 | 5.77 | 7.79 | 4.98 | 6.39 | 7.08 | 6.39 | 7.07 | 10.47 | 5.46 | 6.12 | 7.50 | 6.12 |
|  | R<sup>2</sup> | 0.92 | 0.94 | 0.69 | 0.91 | 0.97 | 0.86 | 0.99 | 0.97 | 0.92 | 0.55 | 0.96 | 0.40 | 0.98 | 0.96 | 0.87 |
| WBAM (% of amplitude) | Bias | -0.0047 | 0.42 | -0.071 | 0.073 | 0.043 | -0.12 | -0.0046 | 0.033 | 0.13 | 0.25 | -0.078 | -0.080 | 0.037 | 0.12 | 0.00018 |
|  | CI | 3.71 | 5.22 | 13.41 | 1.97 | 3.28 | 6.99 | 3.62 | 4.74 | 11.76 | 4.35 | 7.12 | 10.66 | 3.5 | 4.9 | 11.3 |
|  | RMSE | 1.89 | 2.74 | 7.46 | 2.00 | 3.16 | 5.83 | 2.21 | 2.87 | 7.48 | 2.44 | 4.25 | 6.39 | 2.1 | 3.0 | 7.02 |
|  | R<sup>2</sup> | 0.92 | 0.94 | 0.69 | 0.91 | 0.97 | 0.86 | 0.99 | 0.97 | 0.92 | 0.55 | 0.96 | 0.40 | 0.98 | 0.96 | 0.87 |

### CoM

The bias of the CoM ranges from few tenth of mm to a few mm. The CI remains smaller than 2 cm for all tasks, being under 1 cm for all tasks in ML direction. RMSE remains under 1 cm except for **Lean** in AP direction (1.03 cm), being smaller in ML direction for all tasks (Figure 2 A). The coefficients of correlation are excellent (0.98 to 0.99).

**Figure 2:**
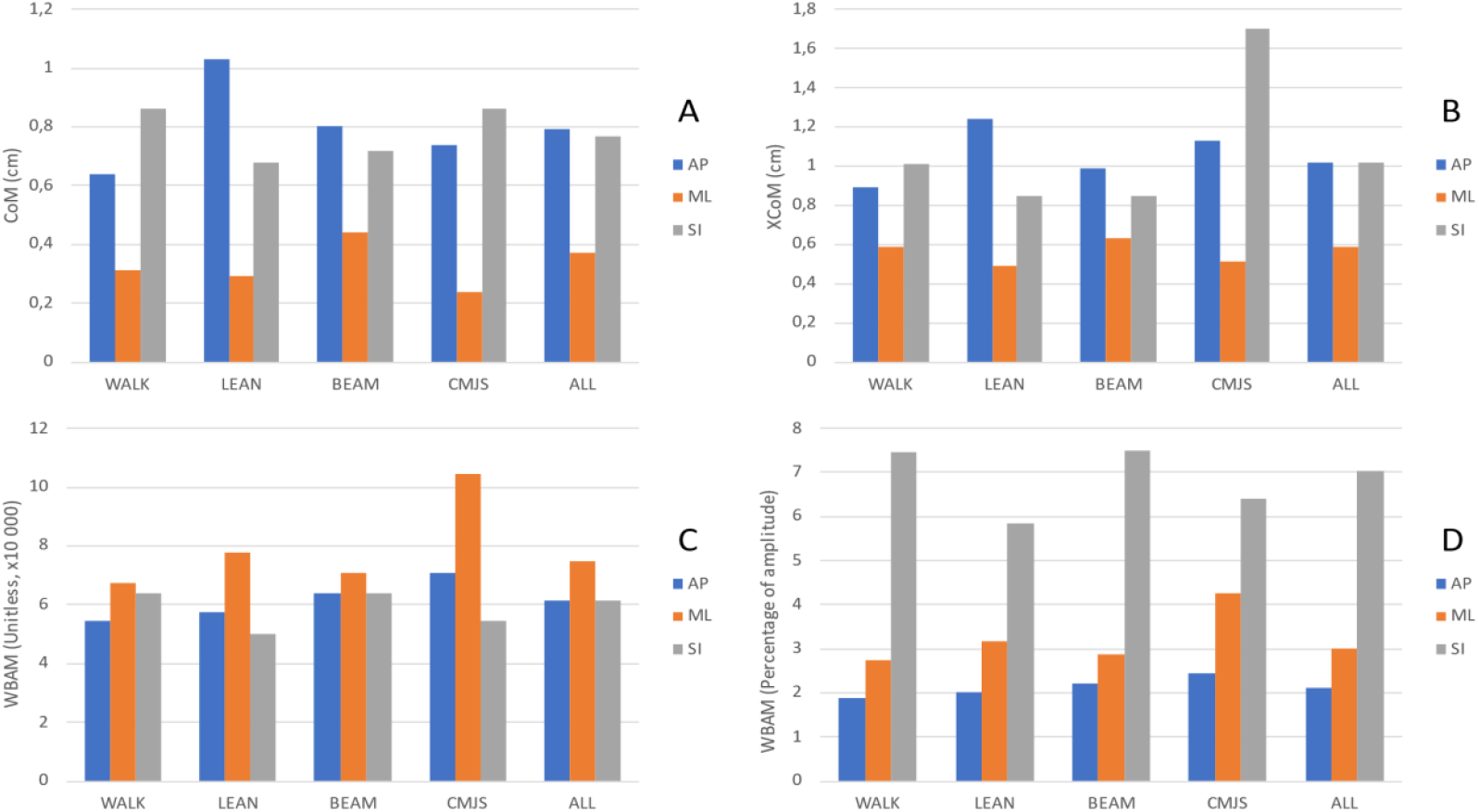
RMSE of the CoM in cm (A), XCoM in cm (B), unitless WBAM (C) and WBAM expressed as a percentage of the amplitude (D), for each task and across all tasks in the AP (blue), ML (orange) and SI (grey) directions.

### XCoM

Regarding the bias of the XCoM across all tasks, its absolute value is bigger in the SI direction, but this is not true for individual tasks except **Walk** and **Beam**. The CI remains smaller than 2 cm except for the **CMJS** in the SI direction (2.76 cm). For both CI and RMSE, the values are always smaller in the ML direction regardless of the task. RMSE mostly remains smaller than 1 cm with few exceptions. Only for the **CMJS** in the SI direction, the RMSE is much higher than 1 cm, but it remains under 2 cm (1.70 cm). The coefficients of determination are excellent (0.97 to 0.99).

### WBAM

The bias across all tasks for the unitless WBAM appears 1000 times smaller in the SI direction than in the AP and ML directions. The order of magnitude of the RMSE of the unitless WBAM across all tasks is the same in all three directions (Figure 2, C), although it does not represent the same percentage of the amplitude (Figure 2, D): the RMSE in the SI direction represents 7.2% of the amplitude of the WBAM, while it is less than 5% in the AP (2.1%) and ML (3.0%) directions. This trend is similar for each task and can also be noticed for the CI. The RMSE expressed in percentage of the amplitude is bigger in the SI direction: the unitless RMSE is of the same order of magnitude as for AP and ML directions but is divided by a smaller amplitude (0.0289, 0.0246 and 0.0085 in AP, ML and SI directions respectively). It thus leads to bigger percentages (Figure 2, C & D). Regarding R^2^, it ranges from poor (0.40) to excellent (0.99), with the **CMJS** being the task with the smallest R^2^ in AP (0.55) and SI (0.40) directions.

## Discussion

This study aimed at assessing commonly used quantities in balance studies, CoM, XCoM and WBAM, estimated with a markerless pose estimation software, Theia3D, and comparing the results to the estimation obtained with a marker-based method. Overall, the magnitude of errors between the two measurement systems is the centimetre for CoM and XCoM and less than 10% of the amplitude for the WBAM.

The RMSE obtained when comparing the CoM computed with the markerless and marker-based systems have a magnitude of around 1cm, and the associated CI remains under 2 cm across all tasks, suggesting a minimal dispersion of the results. Few studies have compared CoM computed using these two approaches. Needham et al [23] focused on computing the CoM during sprinting, using an open source markerless software and a full body marker set. Mean difference of CoM position between the two measurement methods was between 1 mm and 9 mm, depending on the applied filter during post processing. They also provided standard deviations which ranged from 16 mm to 32 mm. The RMSE and CI values obtained in our study for the CoM (from 3.7 mm to 7.9 mm for the RMSE and from 6.8 mm to 15.3 mm for the CI) are consistent with those obtained by Needham et al [23].

The RMSE and CI obtained for the XCoM have the same order of magnitude as the ones obtained for CoM, which is consistent as the XCoM is computed based on CoM. In the literature, the simplified marker set proposed by Tisserand et al [11] has been used to compute the XCoM during gait and fall recovery, and the resulting XCoM was compared to that obtained using the marker set of Dumas et al [33]. Mean distance between the XCoM computed using simplified and reference models ranged from 7.8±3.4 mm to 8.5±2.8 mm. This reported magnitude of error of the XCoM computed using the reference and the simplified models is similar to that obtained in our study. As the simplified marker set was adopted in other studies, this magnitude of error seems acceptable. To the best of our knowledge, no study has been conducted to compare XCoM computed with markerless and marker-based approaches. It is however questionable whether such errors are acceptable when comparing groups with different conditions, especially in the clinical context. In their repeatability study, de Jong et al [34] have computed distances between the XCoM, measured with markers, and the centre of pressure (CoP), measured reliably with an instrumented treadmill, during gait. During one of their measurement sessions, they found differences of approximately 300 mm in the AP direction and 25 mm in the ML direction between the control and patient groups. These differences are more important than the errors observed between markerless and marker-based systems.

However, when comparing the stroke patients and spinal cord injury patient groups, the differences drop to approximately 70 mm and 4 mm in the AP and ML directions respectively, the latest being potentially too small to be detected when using a markerless system as it falls within the CI estimated in our study. It should thus be verified beforehand if the accuracy required for the clinical application is compatible with the errors (bias and CI) that can be found between markerless and marker-based systems.

It is more difficult to compare the results we obtained for the WBAM to the literature because of the different existing normalizations techniques. Begue et al. [32] have studied WBAM, expressed with the same normalization, for young and old adults during stepping at preferred and fast speeds. The smallest significant difference they found was of 0.6×10^−3^ (unitless). It is the same order of magnitude as the RMSE found in our study for the WBAM across all tasks (0.61×10^−3^ to 0.75×10^−3^, unitless). The other differences found in their study are bigger (from 1.6×10^−3^ to 5.9×10^−3^ unitless) and are outside of the CI we found. It seems therefore also possible to use a markerless system to compute WBAM and compare groups and conditions in balance studies, but one should be aware of the order of magnitude of the differences found between the conditions, as the differences could be partly due to using the markerless instead of marker-based motion capture system.

It appears that the task performed and the associated dynamic aspect have an influence on the errors obtained between the two measurement systems, which is consistent with the findings of previous studies [35,36]. When looking at the CoM and XCoM (Figure 2 A & B), the biggest RMSE occurs in the main direction of movement, which is the direction with the most important velocity, for **Lean** (AP direction), **Beam** (AP direction) and **CMJS** (SI direction). The importance of the dynamic aspect is most obvious for the **CMJS** in SI direction: the difference of RMSE between the main direction of movement (SI) and the two other directions is higher for XCoM than for the CoM, as the XCoM includes the velocity of the CoM in its expression (Figure 2 A & B). For the WBAM, the biggest RMSE can also be found around the main direction of rotation for **Lean** task and **CMJS** (both ML direction), although it is less clear for **Walk** and **Beam** tasks. Moreover, for all the tasks, the smallest RMSE of the CoM and XCoM is systematically in the ML direction: as these tasks take place in the sagittal plane, there is little displacement and small velocity in the ML direction, thus the error of measurement is reduced. There is no similar phenomenon for WBAM. This should be considered when studying a task with an important velocity in one direction, as it appears that the biggest error between markerless and marker-based systems can be found in the main direction of movement.

There are some limitations to this study. The first limitation is that participants were healthy young adults, and it is a possibility that the dataset used to train the neural network of the markerless system contained mainly similar subjects. It is thus relevant to wonder if the same results would be obtained with participants with different ages and/or anthropometric characteristics, or patients with body deformities. Moreover, this study was conducted in a controlled laboratory environment: participants were wearing minimally fitting clothes, the lightning conditions were controlled and there was a consequent number of cameras. All these parameters are currently under study in the literature to understand their effect on the accuracy of markerless motion capture [37–39]. Another possible limitation is the different approaches used to build marker-based and markerless models [30], which could generate systematic bias. However, we found very small biases for CoM, XCoM and WBAM, suggesting that the models are consistent.

## Conclusion

In this study, we assessed whether a markerless motion capture system, Theia3D, can be used to compute quantities that are widely used in balance-related studies. The results showed a moderate-to-strong level of agreement between the markerless and marker-based systems. With such errors, a markerless approach thus seems a reasonable alternative to marker-based systems for balance studies.

## Acknowledgements

This study was supported by the ANR grant 20-STHP-0003.

## Notes

### Competing Interest Statement

The authors have declared no competing interest.

